# Yclon: Ultrafast clustering of B cell clones from high-throughput immunoglobulin repertoire sequencing data

**DOI:** 10.1101/2022.02.17.480909

**Authors:** João Gervásio, Alice Ferreira, Liza F. Felicori

## Abstract

**Motivation:** The next-generation sequencing technologies have transformed our understanding of immunoglobulin (Ig) profiles in various immune states. Clonotyping, which groups Ig sequences into B cell clones, is crucial in investigating the diversity of repertoires and changes in antigen exposure. Despite its importance, there is no widely accepted method for clonotyping, and existing methods are computationally intensive for large sequencing datasets.

**Results:** To address this challenge, we introduce YClon, a fast and efficient approach for clonotyping Ig repertoire data. YClon uses a hierarchical clustering approach, similar to other methods, to group Ig sequences into B cell clones in a highly sensitive and specific manner. Notably, our approach outperforms other methods by being more than 30 to 5000 times faster in processing the repertoires analyzed. Astonishingly, YClon can effortlessly handle up to 2 million Ig sequences on a standard laptop computer. This enables in-depth analysis of large and numerous antibody repertoires.

**Availability and implementation:** YClon was implemented in Python3 and is freely available on GitHub

(https://github.com/jao321/YClon.git)

## 1. Introduction

The use of next-generation sequencing in Adaptive Immune Receptor Repertoire (AIRR-Seq) studies provides only a snapshot of the antibody repertoire diversity that is composed of up to 10^18^ distinct antibodies (Briney et al., 2019). Moreover, those advanced techniques help to better understand the immune system as it allows the study of millions to billions of distinct antibody sequences (DeWitt et al., 2016; Briney et al., 2019). To effectively interpret this vast amount of data, a crucial step is the grouping of B cell clones. These clones are derived from a common progenitor B cell (Hershberg et al., 2015) and typically bind to the same epitope. The B cell receptor (BCR) is generated by the rearrangement of V (D) and J gene segments, with the CDR3 located at the junction of these segments playing a critical role in antigen recognition and binding specificity. CDR3 sequence analysis can identify B cell clones with shared antigen specificities, making it a powerful tool for characterizing the B cell repertoire. The size of a B cell clone and the number of clonotypes (each clone is only counted once) can provide insights into repertoire diversity, which is related to different factors such as age (Davydov et al., 2018), response to vaccination or infection (Khavrutskii et al., 2017), allergy (Wu et al., 2014), cancer (Zhang et al., 2019), among other conditions. However, identifying antibody sequences belonging to the same B cell clone remains a challenge (Gupta et al., 2018), particularly with such large datasets.

Some researchers consider the same clone antibody sequences sharing the same V and J genes and identical junction region (CDR3 plus the conserved anchors at positions 104 and 118), as the case of IMGT/HighV-QUEST (Li et al., 2013, Aouinti et al., 2015). In general, there is a consensus that the V and J genes have to be shared by antibodies to be considered as part of the same clonotype. The main difference in the clonotyping methods is the way and the cut-offs used to group the antibodies junction region or the CDR3. Some approaches like the one containing in Change-O (Gupta et al., 2015, Gupta et al., 2017) and SCOPer (Nouri & Kleinstein, 2018, Nouri & Kleinstein, 2020) assigns clones based on the hamming distance of junction, then performs a clustering method (single-linkage for Change-O and spectral clustering with an adaptive threshold for SCOPer). One problem with most of the above approaches is the time to process high-throughput data, with negatively impacts downstream analyses, especially in limited resources laboratories. Lindenbaum et al., 2020 proposed a method to group Ig sequences into clonotypes, that does not require gene assignments and is not restricted to a fixed junction length. What it does is vectorize a sequence of 150 nucleotides, covering the junction, into k-mers of size 5, which should be fast. However, they do not take into account if the sequences are annotated as sharing the same V and J genes. Therefore, it is possible that the clones assigned by Lindenbaum et al, 2020 do not fit the definition of clone hereby established.

The increasing availability of BCR sequencing datasets from different sources presents an enormous potential for in-depth analysis. In order to ensure consistency and quality in reporting this data, the Adaptive Immune Receptor Repertoire (AIRR) Community has proposed the Minimal Information about AIRR (MiAIRR) standard (Rubelt et al., 2017), which outlines the essential information that should be included in reporting these datasets. In this sense, the databases OAS and iReceptor, were created, and gathered, 3,448,993,669 and 815,114,273 IGH sequences, respectively, on May 17th, 2023 (Corrie et al., 2018; Olsen et al., 2021). Notably, 77.7% of IGH sequences in OAS belong to repertoires consisting of over a million sequences, while iReceptors exhibit an even higher percentage at 82%. This shows the need for a more scalable tool, able to clonotype hundreds of antibody repertoire datasets, including large repertoires, quickly and efficiently while respecting the definition of clone.

Because of this, we propose YClon, a tool that rapidly processes repertoires to identify clonotypes, including the ones with more than 2 million sequences, in less than one hour, considering identical V and J genes and similarity in the CDR3.

## 2. Material and Methods

### 2.1. YClon Implementation

YClon is a script written in Python3 and receives as input, on its default setting, “. tsv” files formatted in the MiAIRR data standard, annotated with IgBlast 1.16.0. The file must contain at least four columns, and they should be assigned as “sequence_id” with a unique identifier for the sequence, “v_call” with the V gene annotation, “j_call” with the J gene annotation, and “cdr3” containing the sequence of the CDR3. In order to clonotype the sequences, YClon goes through 5 steps:

#### I. Subsetting

The first step is to subset the data sharing the same V and J gene segments and CDR3 length.

This is achieved using the following steps:

a. We recover v_call, j_call, and cdr3 columns indexes and their associated values in the subsequent rows of the .tsv file.
b. We calculate the length of the cdr3 sequence and the v_call and j_call column’s gene names (without allele information). The combination of these attributes will be the key to a dictionary. For each key, there is a list of tuples containing the sequence_id and the cdr3 sequence.
c. For each new key, a new list is appended to a dictionary. If the key is already in the dictionary another tuple is appended to the list.

#### II. Vectorizing CDR3 nuceotide sequence

CDR3 sequences of the data subsetted were deconstructed in n-grams of size 3 with a sliding window of 1 nucleotide as reviewed by Vinga & Almeida, 2003 and determined using different n-gram sizes as described below (section 2.2).

Yclon can use two different vectorizing methods.

The default method counts the number of times a specific n-gram is detected along the sequence using a sliding window of one nucleotide (Figure 1A) using the CountVectorizer function of the scikit-learn library (Pedregosa *et* al., 2011).

**Figure 1:**
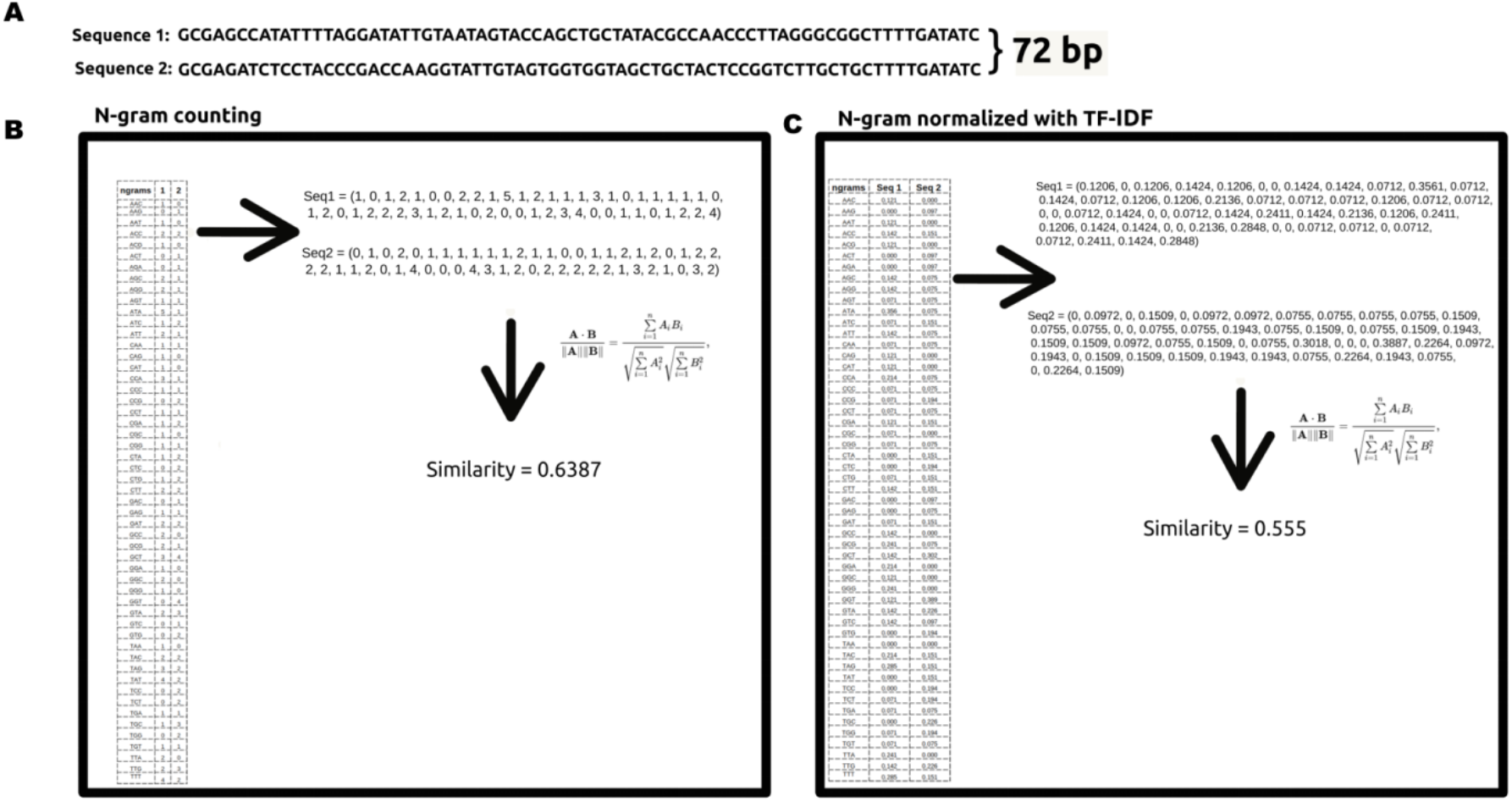
YClon Vectorization Methods: An Overview. This figure illustrates the two vectorization methods used in YClon. (A) Two sequences of equal length, 72 bp, are compared using two different methods described in (B) and (C). (B) The sequences are decomposed into n-grams of size 3 (as determined in this study using simulated repertoires). A vector is formed by counting the n-grams present in the sequence. The cosine similarity between the vectors is calculated to determine the similarity of the sequences. The formula shows how the cosine similarity is calculated between vectors A and B. (C) The sequences are again decomposed into n-grams of size 3. The n-gram frequency is then normalized using TF-IDF (term frequency-inverse document frequency) weighting. This weighting takes into account both the frequency of the n-gram in the sequence and the inverse presence of the n-gram in all sequences being compared. The resulting vector is calculated by dividing the n-gram frequency in the sequence by the total number of n-grams in the sequence (TF) and then multiplying this TF by the inverse frequency (IDF) of the n-gram in all sequences being compared. The IDF is calculated by dividing the number of sequences being compared by the number of sequences containing the n-gram. The cosine similarity between the normalized vectors is calculated to determine the similarity of the sequences.

The user can also choose to use the frequency-inverse document frequency (tf-idf) (Figure 1B) with the TfidfVectorizer function of the scikit-learn library (Pedregosa *et* al., 2011). The tf-idf is calculated in a similar way as the one described in Lindenbaum *et* al, 2020. These methods ignore the locations of particular n-grams within each sequence.

#### III. Comparing vectors

Afterwards, a similarity matrix is built by comparing every vector, representing the CDR3 sequences, using cosine similarity (Sidorov *et* al., 2014).

This is achieved using the cosine_similarity function from the scikit-learn library (Pedregosa *et* al., 2011).

#### IV. Clustering

In the Yclon default method, the resulting square distance matrix is the input for a hierarchical agglomerative method using an AgglomerativeClustering method from the scikit-learn library, with a default threshold distance of 0.09, which was determined as described below (section 2.2) (Pedregosa *et* al., 2011).

In addition to the default method, it is also possible to remove redundant sequences and compare only the unique ones, instead of every sequence from that particular subset. For this, the function drop_duplicates from Phyton3 was used for each key. After the clustering, those collapsed unique sequences are re-expanded using the data of the original key.

#### V. Writing output file

The last step consists of writing an output file containing all assigned clone sequences into the “clone_id” column. If the file already has a “clone_id” column, the new column is called “clone_id_yclon”. The output file will be named by adding “Yclon_clonotyped” to the input file name.

### 2.2. Testing different n-gram sizes and clustering thresholds

We evaluate different n-gram sizes to vectorize CDR3 nucleotide sequences using 74 simulated repertoires data from Lindenbaum *et* al., 2020, by calculating the values of sensitivity, specificity, and PPV.

Specificity [1] was calculated by dividing the number of sequences that were correctly identified as not being in a clone (true negatives or TN) by the total number of sequences that were not in the clone (TN plus false positives or FP).

Sensitivity [2] was calculated by dividing the number of sequences that were correctly identified as being in a clone (true positives or TP) by the total number of sequences that were actually in the clone (TP plus false negatives or FN).

Finally, positive predictive value (PPV) [3] was calculated by dividing the number of true positives (TP) by the total number of predicted positives (TP plus FP). This measure is used to adjust the weight of the PPV in the whole repertoire.

To facilitate the comparison of different thresholds used in the analysis, a new metric called SSP was created by multiplying sensitivity, specificity, and positive predictive value (PPV). The closer the SSP values are to 1, the better the performance of YClon for clonotyping.

To evaluate YClon’s performance, we ran the software using an n-gram size ranging from 1 to 9. We also tested 100 different values to decide the threshold for agglomerative clustering, from 0.00 to 0.99.

To obtain the average CDRH3 length, we accessed iReceptor and filtered their data using the following parameters: Organism as Homo sapiens (NCBITaxon:9096), PCR target as IGH, Cell subset as B cell, germinal center B cell, immature B cell, naive B cell, peripheral blood mononuclear cell, plasmablast. The filtered data were analyzed by measuring the length of every CDR3, multiplying that number by the number of times that CDR3 appeared, and dividing by the total number of sequences.

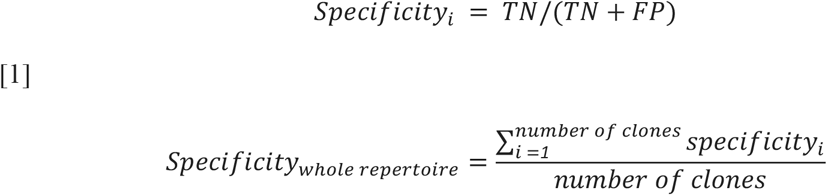

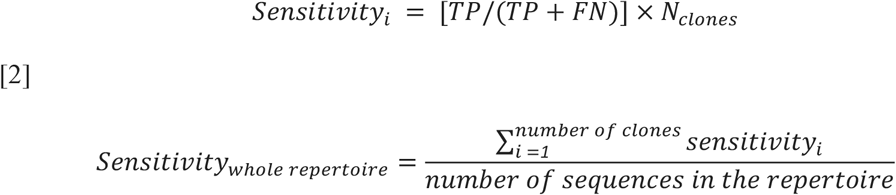

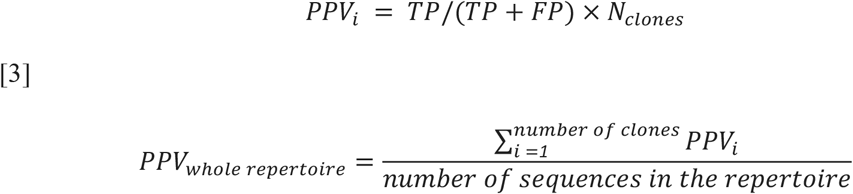

### 2.3. YClon Performance

We tested YClon performance against grounded true data using 74 simulated repertoires, with known clonal relationships, from Lindenbaum et al., 2020.

For this analysis, we also run different clonotyping methods to compare with YClon: Change-O, using thresholds ranging from 0.12 to 0.16 determined using SHazaM (Gupta et al., 2015), SCOPer (Nouri & Kleinstein, 2018) and Lindenbaum’s method (Lindenbaum et al., 2020). The mean values of specificity, sensitivity, and PPV were calculated for each clone and used to assess YClon’s performance.

To evaluate the tool’s functionality, we ran YClon with 3 antibody repertoire datasets of different sizes (Table 1) in a Desktop with an Intel(R) Core(TM) i5-4590 CPU @ 3.30GHz, with 16 Gb of RAM, running ubuntu 20.04 and compared the time each software took to complete the task. The smaller repertoire (Ghraichy et al., 2020) was re-annotated using IMGT/HighV-Quest (Aouinti et al., 2015), while the two others were downloaded already annotated (Chang et al., 2016; Briney et al., 2019) from iReceptor (Corrie et al. 2018) and OAS (Olsen et al., 2021) respectively. We also clonotyped the same repertoires using Change-O, using the threshold of 0.12 determined using SHazaM (Gupta et al., 2015), the hierarchical clustering function of SCOPer (Nouri & Kleinstein, 2018) and the method presented by Lindenbaum et al., 2020 to compare the results with YClon. We also evaluate the upper limit of sequences YClon can handle for each V, J, and CDR3 length combination. For that, we chose a set of 463 sequences that shared the same V, J gene annotations and CDR3 length of a simulated repertoire and run YClon. This procedure was repeated, adding in each cycle 463 more sequences until it reaches the maximum memory capacity and the script stops.

**Table 1:** Details of the datasets of different sizes antibody repertoires used in this study.

|  | **Dataset 1** | **Dataset 2** | **Dataset 3** |
| --- | --- | --- | --- |
| **Number of sequences** | 82,966 | 365,370 | 1,741,413 |
| **Database containg Sequences** | OAS | iReceptor | iReceptor |
| **Study ID in iReceptor** | - | PRJNA628125 | PRJEB9332 |
| **Subject** | C.037.1 | 7450 | Family 4 Carrier |
| **Sample** | - | M369-S001 | F4Cpost |
| **Study** | Ghraichy *et al.*, 2020 | Nielsen *et al.*, 2020 | Chang *et al.*, 2016 |
| **Sex** | Male | Female | Female |
| **Age** | 4 | 73 | 3 |
| **Condition** | Healthy | Covid-19 | hepatitis vaccination, 1 day after |

### 2.4. CDR3 intraclonal similarity

To assess the intraclonal similarity of CDR3 sequences generated by YClon, Change-O, SCOPer, and Lindenbaum’s methods, we used the hamming distance to compare the sequences within each clone. We applied this calculation to the 800 largest clones from the 74 repertoires described in Lindenbaum et al., 2020. We used the python library “distance” and the function “hamming()” to compute the proportion of differences between each pair of sequences, assuming they were aligned due to equal length. The average similarity was calculated for each clone.

## 3. Results and Discussion

In this study, the optimal n-gram size and cut-off for vectorizing and clustering sequences were determined using 74 simulated antibody repertoires. The results indicated that an n-gram size of 3 (Figure 2A) and a threshold value of 0.09 (Figure 2B) yielded the highest SSP score, making them the default settings implemented in YClon.

**Figure 2:**
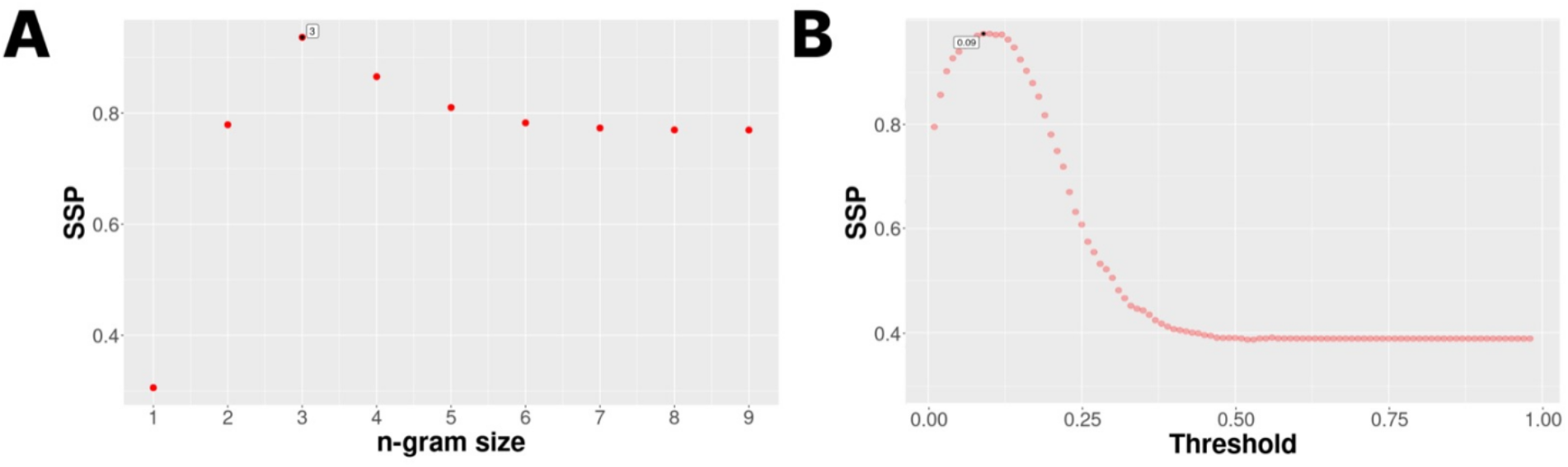
Evaluation of YClon using Different N-gram Sizes and Clustering Thresholds. To verify the best Yclon n-gram size and threshold we clonotyped 74 simulated repertoire and calculated the values of sensitivity, specificity, and positive predictive value (PPV). We combined these values into a single metric, SSP (sensitivity x specificity x PPV), which ranges from 0 to 1, with a higher value indicating better clonotyping results. (A) Evaluation of YClon using n-gram sizes ranging from 1 to 9 bases. (B) Evaluation of YClon using threshold values ranging from 0 to 1, with increments of 0.01.

In contrast, Lindenbaum et al. (2020) utilized an n-gram size of 5 for comparing 150-nucleotide sequences, specifically targeting the junction region. In our approach, we prioritize sensitivity, specificity, and positive predictive value (PPV) rather than solely focusing on sensitivity. Additionally, we only consider sequences with the same length of complementarity-determining region 3 (CDR3), which is shorter than the region examined by Lindenbaum et al. The average CDR3 length in the iReceptor dataset, which consisted of 122,229,351 sequences, was found to be 47.52 nucleotides (equivalent to 15.90 amino acids). This length is less than one-third of the sequence length employed by Lindenbaum et al. These differences in sequence length and region targeted may explain why YClon achieved better results with a smaller n-gram size.

YClon utilizes a fixed threshold of 0.09 to group sequences into clones based on a simulated antibody repertoire of 74. In contrast, other clonotyping methods like SCOPer dynamically adjust the threshold during clonotyping, Change-O recommends using SHazaM to determine the optimal threshold per sample, and Lindenbaum’s method defines the threshold using a negation table. The use of a fixed threshold offers advantages such as simplicity, reproducibility, standardization, and reduced computational complexity. However, if users wish to modify the threshold, they have the flexibility to run SHazaM (Gupta et al., 2015) or other methods to predict a specific repertoire threshold. For instance, in the case of the simulated repertoire, a threshold of 0.12 was found using SHazaM, which closely aligns with the threshold identified by YClon.

Upon examining the intraclonal CDR3 similarity across different clonotyping methods, we observed that YClon, Change-O, SCOPer, and Lindenbaum’s methods primarily grouped sequences within clones exhibiting over 95% similarity (Figure 3). Nevertheless, a small number of outliers with lower sequence similarity were identified in each method, ranging from 45.2% to 94.2% identity among sequences within the same clone. These outliers accounted for 7.88% of the sequences in SCOPer, 3.29% in Change-O, 5.36% in YClon, and 3.83% in Lindenbaum’s method (Figure 3).

**Figure 3:**
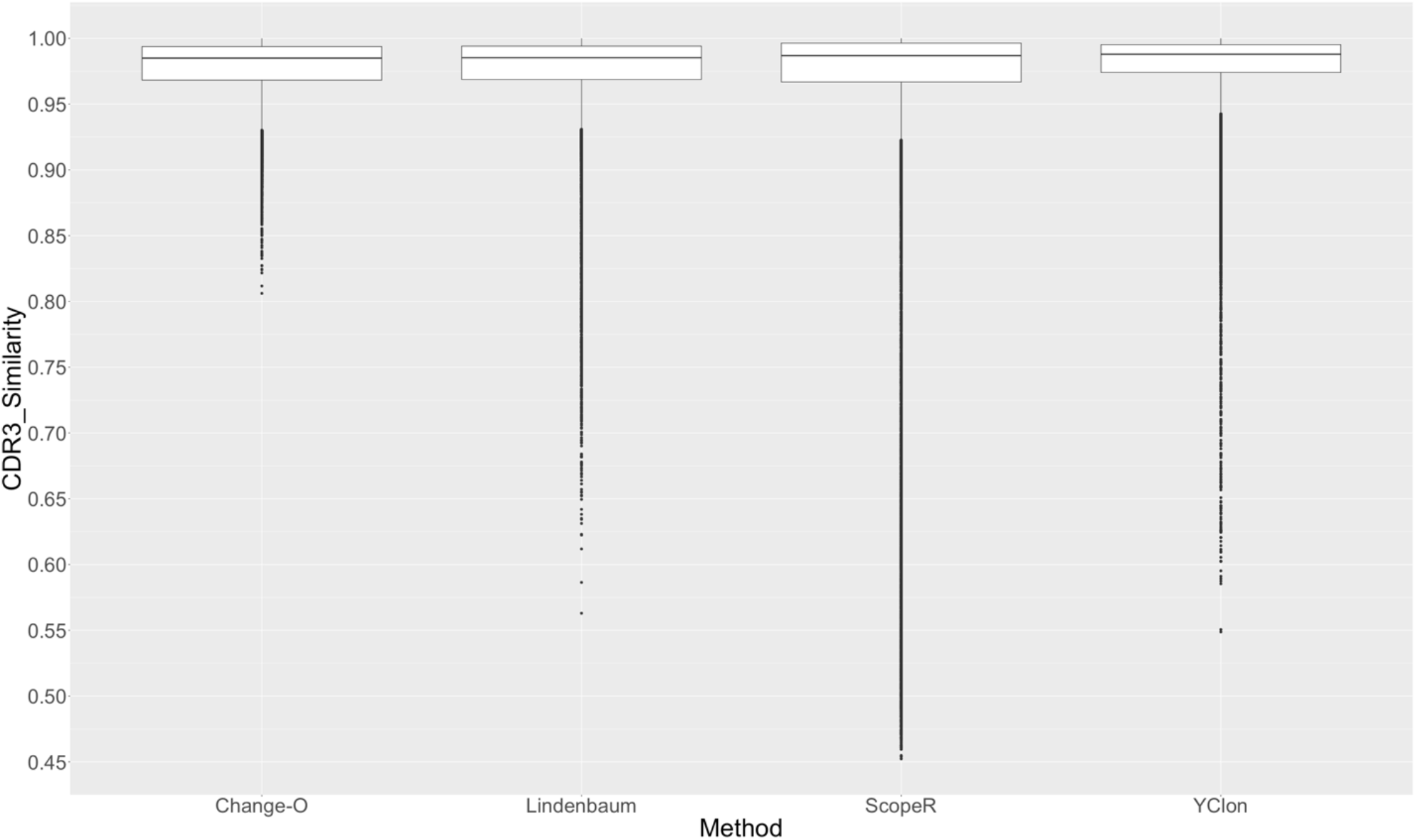
Sequence similarity of the CDR3 sequences within each clone. We applied this calculation to the 800 largest clones from the 74 repertoires described in Lindenbaum et al., 2020. The box plots enclose the 25th-100th percentiles of the data, with whiskers extending to the 25th percentiles and individual outliers represented by circles. Outliers were defined as data points beyond 1.5 times the interquartile range from the lower and upper hinges of the box plot. The metrics used to define the intraclonal similarity is explained in the 2.4 section.

We conducted an assessment to determine the maximum number of sequences that YClon can process simultaneously for each V, J, and CDR3 length combination. Our findings revealed that YClon can compare a maximum of 93,526 sequences in a single run.

To evaluate the performance of YClon, we utilized a dataset consisting of 74 simulated antibody repertoires, where the correct clonal assignments were known. This enabled us to calculate key metrics such as sensitivity, specificity, and positive predictive values (PPV) for YClon, allowing for a comprehensive comparison with other clonotyping methods, including Change-O, SCOPer, and Lindenbaum’s method.

All methods demonstrated high specificity values, approximately 100% (Figure 4A), indicating that the majority of sequences were correctly assigned to the same clone by the respective methods. In terms of sensitivity, YClon exhibited the best performance (96.4%) compared to SCOPer (93.6%), but it was surpassed by Change-O (99.9%) and Lindenbaum’s method (97.18%) (Figure 4B). Regarding positive predictive value (PPV), YClon achieved a higher value (99.8%) compared to SCOPer and Lindenbaum’s method (96.7%), and a similar value to Change-O (99.7%) (Figure 4C). It is important to acknowledge that these results may be influenced by the specific characteristics of the simulated repertoire used for evaluation purposes. It is noteworthy that the results obtained in the SCOPer paper (Nouri and Kleinstein, 2018) align closely with our findings in terms of specificity and sensitivity. However, there is a discrepancy in the node positive predictive value (PPV) calculation compared to our study, where Lindenbaum et al. (2020) reported an 84.7% PPV, whereas we observed a value exceeding 90%. This divergence might stem from variations in the calculation methodologies employed for these parameters. On the other hand, the Change-O paper (Gupta et al., 2015) did not include any data pertaining to sensitivity, specificity, or PPV. Consequently, it is not feasible to make a direct comparison of results with the Change-O paper. Overall, Change-O achieved a higher sensitivity-specificity-positive predictive value (SSP) value (99%) compared to YClon (96%), SCOper (90%), and Lindenbaum’s (82%) methods (Figure 4D). However, when evaluating the performance of each method using 3 antibody repertoire datasets of different sizes (presented in Table 1), YClon showed improved performance, with a clonotyping speed that was 33 times faster than SCOPer and 82 times faster than Change-O for the smaller repertoire (11s versus 6m13s versus 15m12s)(Table 2). For the medium-sized repertoire, Yclon was 301 times faster (58s) than SCOPer (4h51s) and 73 times faster (1h11m20s) than Change-O (Table 2). For a large dataset containing over 2 million sequences, YClon was the only method that could run on the computer, taking 15 minutes and 44 seconds (Table 2). Lindenbaum’s method took the longest time to complete the clonotyping process. Yclon was more than 5000 faster to clonotype medium-size repertoire compared to Lindenbaum’s method (58s vs 16h19m). In summary, YClon not only achieved competitive accuracy in clonotyping, but it also excelled in terms of speed and scalability, making it a highly efficient and practical choice compared to Change-O, SCOPer, and Lindenbaum’s method.

**Table 2:** Performance Comparison of YClon, Lindenbaum’s Method, SCOPer and Change-O for Clonotype Identification across Different Repertoire Sizes.

|  | Dataset 1 |  |  |  | Dataset 2 |  |  |  | Dataset 3 * |
| --- | --- | --- | --- | --- | --- | --- | --- | --- | --- |
|  | YClon | Lindenbaum | SCOPer | Change-O | YClon | Lindenbaum | SCOPer | Change-O | Yclon |
| Number of sequences | 82,928 |  |  |  | 365,707 |  |  |  | 2,081,722 |
| Total number of clones | 24,127 | 978 | 22,680 | 24,031 | 73,741 | 27,921 | 60,561 | 72,446 | 1,610,373 |
| Clones with more than 1 sequence | 15,833 | 673 | 14,477 | 15,648 | 35,537 | 9,783 | 15,095 | 35,474 | 203,766 |
| Benchmark time | **11s** | 30m54s | 6m13s | 15m12s | **58s** | 16h19m | 4h51m | 1h11m 20s | **15m44s** |
| Number of clones that represents the most abundant 5% sequences | 35 | 1 | 23 | 33 | 2 | 1 | 2 | 2 | 946 |
| Number of clones that represents the most abundant 15% sequences | 376 | 1 | 162 | 327 | 22 | 2 | 15 | 19 | 32,836 |
| Number of clones that represents the most abundant 25% sequences | 1,180 | 1 | 719 | 1,190 | 305 | 3 | 94 | 276 | 126,468 |

\* For the dataset 3 it was only possible to run YClon in the computer used for clonotyping. The abbreviations "s" refer to seconds, "m" to minutes, and "h" to hours.

**Figure 4:**
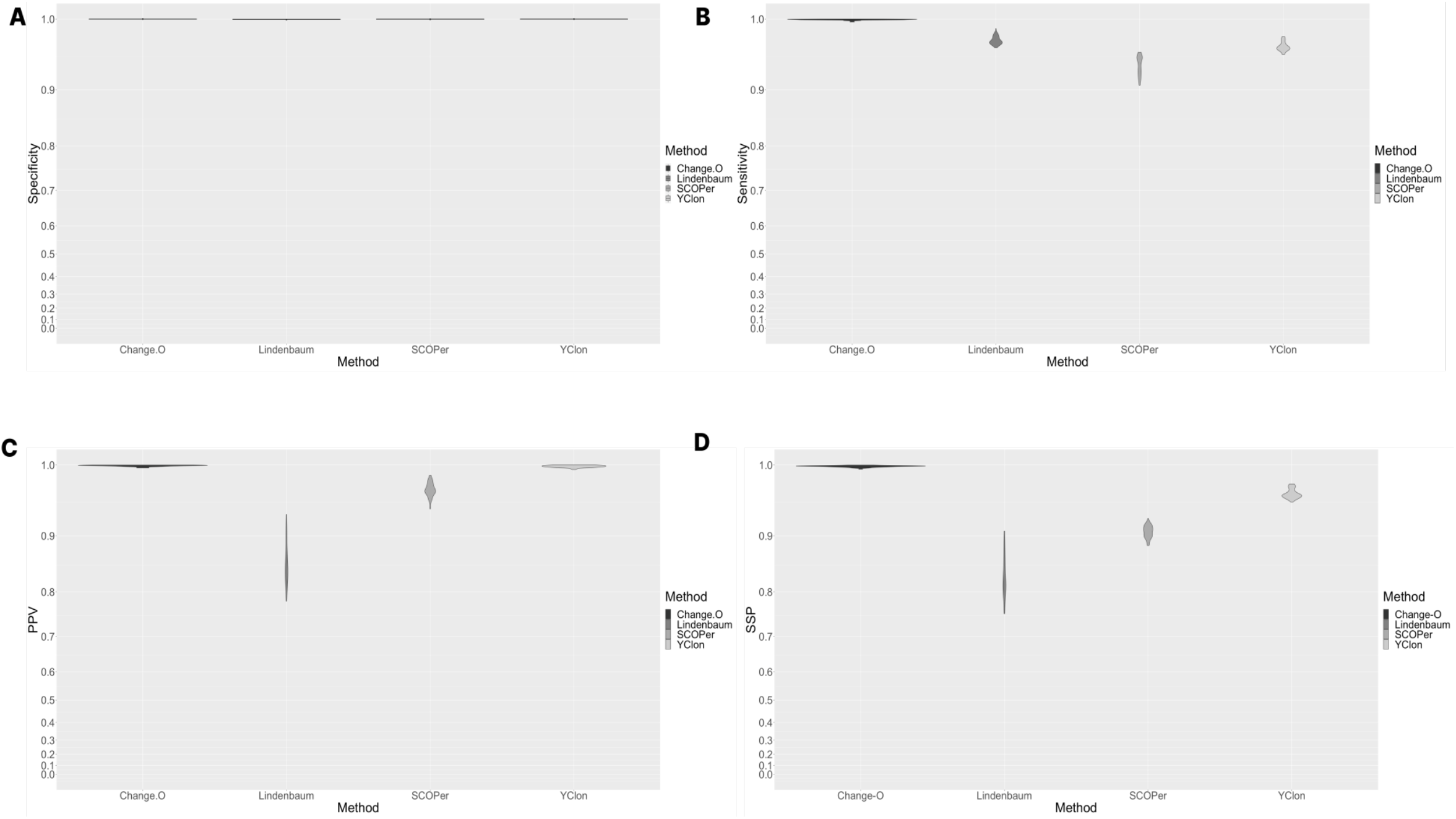
Simulated repertoire clone identification metrics. Violin plot displaying the values for (A) specificity, (B) sensitivity, (C) positive predictive value (PPV), and (D) SSP (sensitivity x specificity x PPV) in clone identification of 74 simulated repertoires from Lindenbaum et al. (2020). The specificity (A) indicates the ability to identify sequences that are not part of a clone, while sensitivity (B) indicates the ability to identify sequences that are part of a clone. PPV (C) represents the probability that sequences in a clone are indeed part of it. The analysis was performed using four clonotyping methods: YClon, the method presented here, Change-O, SCOPer, and Lindenbaum’s method.

## Conclusion

In recent years, the demand for the analysis of large antibody repertoire datasets, containing millions of sequences, has been on the rise. The iReceptor Gateway integrates over 8,919 antibody repertoires, with 856 of them having more than 1 million sequences, and OAS has 15,148 repertoires with 631 of them having more than 1 million unique sequences. YClon provides a solution to this challenge by making clonal assignments faster, more reliable, and possible on an ordinary computer. By surpassing the computational constraints, YClon enables researchers to acquire valuable insights from antibody repertoire data, thereby facilitating the progression of our comprehension regarding the immune system.

## Authors Contribution

JG designed the study, performed analysis, generated the data and wrote the paper. AF analyzed and generated data. LFF designed the study, discussed the results and wrote the paper.

## Acknowlegdement

Coordenação de Aperfeiçoamento de Pessoal de Nível Superior – Brazil (CAPES) [schollarship to JG, grant numbers 88887.506611/2020-00, 88887.504420/2020-00 and 935/19 (COFECUB)]; Fundação de Amparo a Pesquisa de Minas Gerais (FAPEMIG) [grant numbers PPM-00615-18, Rede Mineira de Imunobiologicos grant #REDE-00140-16]; Conselho Nacional de Desenvolvimento Científico e Tecnológico (CNPq) [Pq to LFF] and through the Instituto Nacional de Ciência e Tecnologia em Venenos e Antivenenos (INCT-INOVATOX, grant no. 406816/2022-0); National Institutes of Health (NIH) [grant number 1R01AI143552-02]; Pro-Reitoria de Pesquisa da Universidade Federal de Minas Gerais. We thank also ChatGPT Jan 30 Version for some english sentence improvement.

